# Understanding PCR Processes to Draw Meaningful Conclusions from Environmental DNA Studies

**DOI:** 10.1101/660530

**Authors:** Ryan P. Kelly, Andrew Olaf Shelton, Ramón Gallego

## Abstract

As environmental DNA (eDNA) studies have grown in popularity for use in ecological applications, it has become clear that their results differ in significant ways from those of traditional, non-PCR-based surveys. In general, eDNA studies that rely on amplicon sequencing may detect hundreds of species present in a sampled environment, but the resulting species composition can be idiosyncratic, reflecting species’ true biomass abundances poorly or not at all. Here, we use a set of simulations to develop a mechanistic understanding of the processes leading to the kinds of results common in mixed-template PCR-based (metabarcoding) studies. In particular, we focus on the effects of PCR cycle number and primer amplification efficiency on the results of diversity metrics in sequencing studies. We then show that proportional indices of amplicon reads capture trends in taxon biomass with high accuracy, particularly where amplification efficiency is high (median correlation up to 0.97). Our results explain much of the observed behavior of PCR-based studies, and lead to recommendations for best practices in the field.

## 1 Introduction

Surveying the natural world by amplifying and sequencing DNA from environmental sources such as water, air, or soil has long been commonplace in microbial ecology [see, e.g., 1, 2, 3], but has recently become popular for characterizing ecological communities of eukaryotes [4, 5, 6, 7, 8, 9]. Because the source of samples is the environment itself rather than specific target organisms, the data resulting from such studies have become known as environmental DNA (eDNA) [8]; the ultimate source of genetic material in the environment may be living or waste cells or extracellular DNA [8]. Techniques that take advantage of such data may include non-PCR-based methods such as hybridization, but generally include an amplification step such as quantitative PCR, digital-droplet PCR, or traditional PCR from mixed templates followed by high-throughput sequencing. This last technique is known as metabarcoding, eDNA amplicon-sequencing, or more generally, marker-gene analysis.

Patterns of diversity have been a focus of metabarcoding studies [10, 11], but in many cases, results from eDNA sequencing may differ substantively from results from traditional, non-PCR-based biodiversity surveys [12, 13, 14, 15]. To evaluate metabarcoding as a tool for assessing biodiversity, we provide a mechanistic, simulation-based approach to understanding the processes that lead ultimately to metabarcoding data.

Ecological inquiry often begins with uncovering patterns of biodiversity, yet sampling biodiversity is inherently difficult and the methods highly varied. Methods for surveys of fish diversity differ fundamentally from surveys of birds or trees. Every way of surveying the world has a different set of processes intervening between the sampled phenomenon (say, the number of different types of snails on a rock) and the recorded observation (the number of snails recorded, trapped, or otherwise counted). The reason that different survey techniques offer different results and insights – even for the same survey target – is because these intervening processes differ between techniques. Some methods have only trivial intervening processes: counts of snails on a rock are subject to ascertainment bias and sampling error, but we expect these counts to be reasonably direct reflections of the “truth” that exists in the world. Environmental DNA provides the potential for standardizing of sampling among disparate species groups – for example, a single sampled bottle of ocean water can be used to survey fish, plankton, benthic invertebrates and mammals. However, producing biodiversity estimates from eDNA sequences requires complex laboratory processes – from collection to extraction through amplification and sequencing – that may substantially affect estimates of biodiversity derived from eDNA.

Specifically, eDNA methods often use PCR, which causes two key differences from other sampling methods. First, PCR exponentially increases the very low concentrations of DNA collected in the environment to make amounts sufficient for further analysis. This exponential process means that stochasticity and small biases in the PCR process can lead to large differences the abundance of each species’ amplicons relative to DNA concentrations in the field [16, 17, 18]. The issues surrounding amplification bias in mixed-template PCRs have long been documented [19, 20], and in the metabarcoding context have recently come under useful scrutiny [21, 22, 23]. Compounding the bias problem is a crucial second difference between PCR-based methods and others: DNA from different species often amplifies at different rates, such that each PCR cycle preferentially amplifies templates with greater affinity for the primers being used (*i.e*., amplification bias) [24, 25]. Furthermore, in contrast to many traditional sampling techniques, metabarcoding datasets are compositional [26]: their information content has an “arbitrary total imposed by the instrument” [26], which necessarily means amplicon counts are not directly related to counts of template molecules in the sampled environment. Many PCR-based analyses of ecological communities gloss over potential biases that arise from using genetic methodologies, and few attempt to quantify either the degree of this bias or its effects on study results. However, understanding the results of metabarcoding surveys requires that we understand how these processes influence estimates of diversity and other survey outcomes.

Here, we briefly review the processes involved in metabarcoding surveys. We then simulate sets of biological communities and subject them to simulated PCR-based processing. We independently vary the important axes of variation for eDNA surveys – specifically the number of PCR cycles and the distribution of taxon-specific amplification efficiencies – to illustrate the effects of these parameters on estimates of biodiversity. We base these simulations on real-world use-cases, parameterizing our models using empirical data where possible. We then evaluate the quantitative performance of taxon-specific amplicon-abundance indices vs. biomass in simulations, finding that proportional indices of eDNA reads capture trends in taxon biomass with high accuracy, particularly where amplification efficiency is high. Our results explain much of the observed behavior of PCR-based studies, and lead to recommendations for best practices in the field.

## 2 Methods

### Major Processes Involved in Metabarcoding

At least five major processes drive patterns of metabarcoding data and affect estimates of biodiversity. Genetic material sampled from an environment derives from some living species (*process 1*). For single-celled species, an organism and its representative genome are coincident, while for multicellular species the sampled DNA may derive from a residual or waste cell (or a gamete) in the environment. DNA presence in the environment – having been created by the source organism and not yet degraded or lost – is then the first process with which we are concerned. In either the single- or multicellular case, the time-averaged DNA shed into the environment is proportional to the biomass of a given species, although the proportionality constant may vary between species [CITE degradation studies]. This DNA degrades rapidly in ambient conditions [27, 28] and may be transported away from the source organism [29, 30], taken up by other organisms via transformation [31], or adsorbed onto soil or other substrates [32]; these mechanisms may be treated together as forms of effective eDNA loss. For a fixed population in a closed area, we hypothesize the observable DNA concentration will be an equilibrium between the generation and loss functions; our simulations assume equilibrium in order to model eDNA template concentration as a point estimate.

This DNA is then sampled by a researcher (*process 2*), extracted and purified from its surrounding cellular matrix (*process 3*), and subject to PCR amplification (*process 4*). This amplification step is of special importance, since it is what most obviously distinguishes genetic sampling methods from traditional ecological sampling. In some applications, a sample is subjected to multiple PCR processes, but at minimum, the amplicons are sequenced (*process 5*) before bioinformatic analysis. Because these (minimum) five processes occur in series, random and systematic errors at one step propagate through the analytical chain [33]. It is therefore important to understand each process individually so that we can estimate their cumulative effects on measures of diversity.

Defining the target community is an important *a priori* component of all studies of biodiversity. While this is widely appreciated in the ecological literature, it is often overlooked in metabarcoding studies. For example, ecologists might study the biodiversity of forest trees [34] or coral reef fish [35] or sessile invertebrates [36]. In the metabarcoding context, very specific primer sets targeting a relatively small number of taxa (e.g., vertebrates [37, 38]) may have a well-defined target group, but nevertheless the absence of a taxon from a sequenced sample does not indicate the absence of that taxon from the environment. Instead, the unsampled species simply may not have been susceptible to that set of PCR primers, and so failed to amplify. The result is often a dataset that represents many taxa, but these taxa are an unknown fraction of a larger (and perhaps spatially or taxonomically undefined) pool of species present. Here, for clarity of illustration, we treat the 1000 simulated species as the eukaryotic community of a nearshore marine habitat, but we note these simulations are broadly applicable to most ecosystems in which PCR-based studies occur.

### Community Simulations

To test the effect of eDNA processing on estimates of abundance and biodiversity, we simulated biological communities and performed simulations of metabarcoding processes on each, as described below.

#### Biomass in the environment (*process 1*)

We generate three different distributions of biomass proportions to test for an effect of these underlying community distributions on metabarcoding diversity estimates (Figure 1). Let *B_i_* be the proportional biomass of species *i*, for *i* = 1,…,*N* species such that 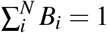. First, we simulate a community in which all species have identical proportional biomass,

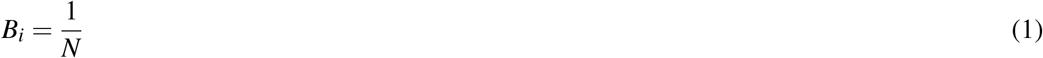

and refer to this as our “uniform” community. The two other communities are defined using a symmetric Dirichlet distribution to describe communities with variation in biomass among species,

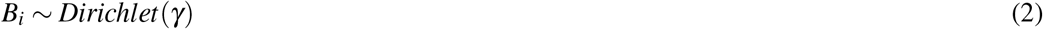

**Figure 1.**
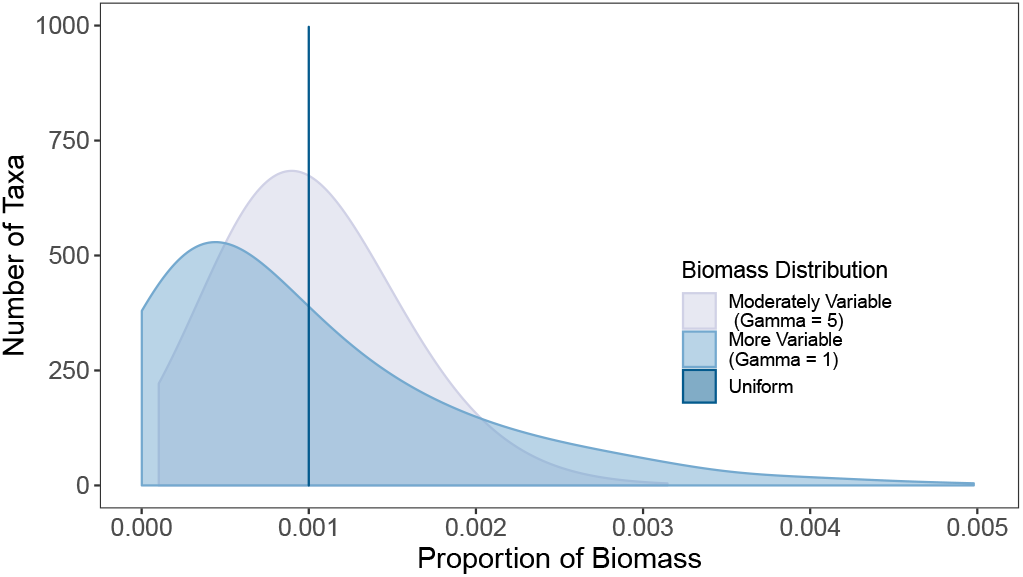
Distribution of proportional biomass in the three types of ecological communities simulated.

We define *γ* = 5 for one community and *γ* = 1 for the second; smaller values of *γ* correspond to more variation among species in proportional biomass. Across all three of these community distributions, the number of species is consistent, as is the expected proportion of each species (i.e., the mean). These characteristics facilitate comparisons across distributions by reducing the differences under consideration to one dimension. We note in particular that in using proportional (rather than absolute) biomass, we can flexibly capture changes in community structure appropriate to those that genetic assays likely respond to: because PCR is a competitive reaction among template molecules, absolute biomass (and hence absolute DNA concentrations) are less relevant than their proportions in the sampled community. We simulate 100 independent communities of 1000 taxa for each community biomass distribution.

#### Genetic Material in the Environment (eDNA, *process 2*)

We assume that organisms shed DNA into the environment, *D_i_*, as a function of the biomass of species *i* times the shedding rate of that species, *s_i_*. While loss of DNA from the environment plays a vital role determining the equilibrium DNA concentration in the environment, we assume that loss of eDNA from the environment is constant among species, and therefore equilibrium DNA concentration is proportional to DNA shedding. Here, we inform our shedding rate parameters using Sassoubre et al. 2016 [28], which found shedding rates (in pg/hour) among three Pacific fishes to vary by two orders of magnitude; accordingly we sample simulated shedding rates from a distribution with wide variance and a moderate central tendency.

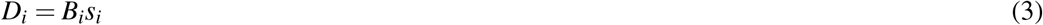

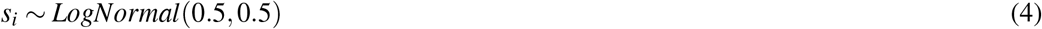

We note that this simulation is spatially inexplicit, and so the statistical distributions of biomass and eDNA are not intended to reflect a particular spatial distribution.

#### DNA Collection and Extraction (*process 3*)

We assume DNA is collected in proportion to its abundance in the environment and extracted with equal efficiency from all species present.

#### DNA Amplification during PCR (*process 4*)

Because each taxon (or, more broadly, template molecule) has its own amplification efficiency for a given set of primers, we describe three simulations according to the distribution of these efficiency parameters among the taxa in a community (Figure 2). We treat biases arising from sequence variation or from secondary structure (“polymerase bias”; see e.g., [39]) as equivalent for the present purposes. For all scenarios, we use the same relationship for translating among-taxon variation in amplification efficiency into the number of amplicons observed for taxon *i, A_i_*, at the conclusion of the PCR. Let *a_i_* be the binding affinity for the PCR primers to species *i*, and *N_PCR_* be the number of PCR cycles, then

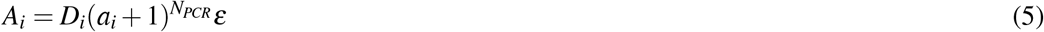

with *ε* representing a multiplicative process error term which adds a small amount of stochasticity to the observed amplicons for each species. We model *ε* as a lognormal distribution, *ε* ~ *LogNormal*(*μ, σ*^2^) with *μ* = 0 and *σ* = 0.05.

**Figure 2.**
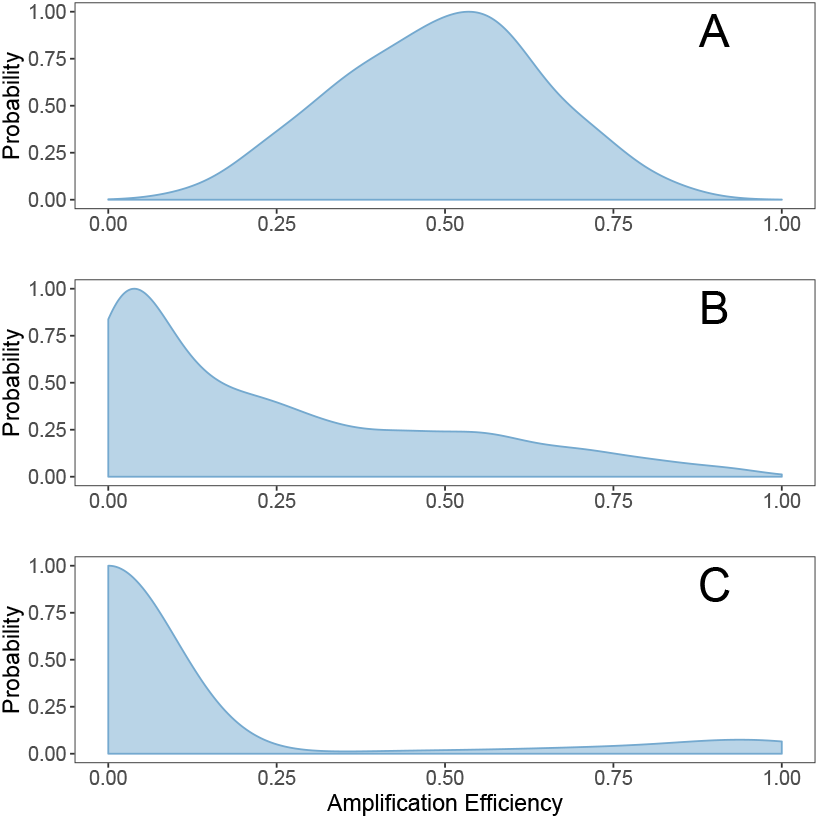
The distribution of amplification efficiencies for three eDNA use-cases. **A**: amplifies most taxa, but few very well or very poorly (symmetrical; likely reflects the performance of broad-spectrum primers such as [44] and [45] acting upon eukaryotes); **B**: amplifies few taxa well, most taxa poorly (right-skewed; parameterized based upon [11]; see main text); **C**: amplifies a small number of taxa very well, but most not at all (parameterized based upon the performance of 12s primers [37] in detecting fish assemblages as described in [41]; see main text)). Note that qPCR primers are a special case of this last distribution.

Key parameters governing the observed number of amplicons from a given eDNA sample are the DNA concentration, *D_i_*, and the amplification efficiency for each species, *a_i_*. For a given eDNA sample, *D_i_* is constant, so we focus on three distributions of amplification efficiencies corresponding to biological use-cases. For each case, we model the amplification efficiency for each species as a draw from a beta distribution,

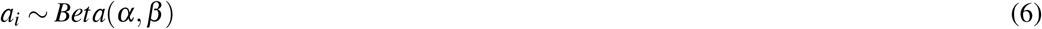

or as a mixture of two Beta distributions,

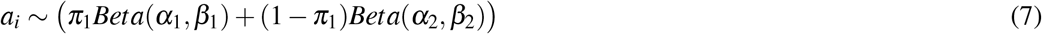

Where *π*_1_ is the weight for the first mixture component and so 0 < *π*_1_ < 1.

We use a mixture of only two distributions, but future work could consider mixtures with larger number of component distributions (for example, where different taxonomic groups make up different components of the mixture). The parameter *a_i_* is equivalent to the measure of amplification efficiency, *E* that is often reported for an individual species in qPCR studies [40].

Any primer set will be evaluated relative to its intended target set of taxa. Put differently, the way we think about primer efficiency depends strongly upon taxonomic scale. Primers designed to amplify vertebrates exclusively might behave very well (i.e., have relative amplification efficiencies clustered around one) within Vertebrata, but across the tree of life, vertebrates are a vanishingly small fraction of biodiversity. Accordingly, at the scale of the tree of life, these same primers would have efficiencies clustered near zero (they do not amplify most lifeforms at all) with a small proportion of target molecules (vertebrate species) amplifying quite well. For simplicity and ease of comparison, we evaluate our simulated primers on a common taxonomic scale, Eukaryota.

We drew empirical data from published metabarcoding papers to parameterize our models, finding several recent papers [41, 4, 11, 42, 43, 39] that reported results from mock (i.e., synthetic) eDNA communities useful for our purposes. These papers provided the number of PCR cycles used, the starting concentrations of DNA for a variety of taxa, the primers used, and the ending counts of amplicon reads; such data allowed us to calculate taxon-specific amplification efficiencies for each primer set (Supp. Table 1). We estimated the parameters for a univariate beta distribution to the observations for each primer, and used the beta parameters to inform our simulations (see below; Figure 2).

##### Case A: Amplifies Most Taxa, but Few Very Well or Very Poorly

For eukaryotes, several primers have been widely used in metabarcoding studies because they amplify eukaryotic taxa across many domains of life (Leray COI primers [44] or the Stoeck 18S primers [45]). It is not yet clear what the distribution of amplification efficiencies is for these primers across Eukaryota, but given the breadth of observed taxonomic coverage (e.g., [46, 12]) here we model these efficiencies using a beta distribution (*α* = 5, *β* = 5), with a mean of 0.5 and a standard deviation of 0.15.

##### Case B: Amplifies Few Taxa Well, Most Taxa Poorly

If we envision the sampled community as being made up of one thousand eukaryotic species, the primers developed for broad-spectrum use and widely useful for population genetics are likely to have a right-skewed distribution when viewed at the scale of Eukaryota. We model this as a beta distribution (*α* = 0.5, *β* = 1.5) with mean 0.25 and standard deviation 0.26. For example, metazoan 16S primers developed in Kelly et al. 2016 [47] amplify very poorly the single-celled photosynthesizers that comprise the majority of eukaryotic DNA in marine environments. Instead, this primer set amplifies animal DNA well and almost exclusively; the result will be a right-skewed distribution (i.e., a mode near zero with a long tail in the positive direction) at the scale of Eukaryota.

##### Case C: Amplifies a single taxonomic group well, most taxa poorly or not at all

The third use-case is analogous to specialized primers used in taxon-specific metabarcoding studies, such as those targeting vertebrates specifically [37, 4, 48, 38]. These target a narrow range of species for a particular survey purpose, and consequently amplify a very small fraction of eukaryotic life present in most environments. For the marine environment, we envision a primer that amplifies vertebrate species (e.g., fish and marine mammals) well but amplifies non-vertebrate taxa little or not at all. We note that qPCR primers are an extreme case of this distribution, in which the primer set exclusively amplifies a single taxon. We model this as a mixture of two distributions: one for the target taxon, and one for other eukaryotes present in the environment. Using parameters derived from [41] (see Supplementary Table 1), we model the target-taxon component as a Beta distribution (*α* = 2.1, *β* = 0.58; 10% of the taxa present), and non-target component as Beta (*α* = 0.01, *β* = 10; 90% of the taxa present).

#### DNA Sequencing (*process 5*)

Finally, the number of sequencing reads for species *i, Y_i_*, is proportional to *A_i_*. The resulting community of eDNA reads is a Multinomial sample of between 10^5^ and 10^6^ reads out of a total of 10^7^ reads – the size of an average Illumina MiSeq run – from the community of amplicons present. The result is a set of replicate samples that varies in read-depth, consistent with common outcomes of MiSeq (and similar) sequencing runs.

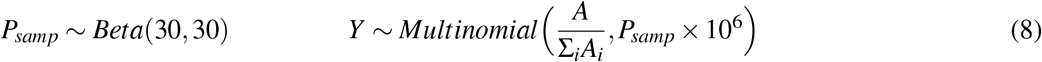

where *Y* is a vector containing the observed amplicon counts for the 1000 species.

### Analyses of Simulations

We used the simulation results for eDNA to understand the characteristics of eDNA data with respect to two important areas of ecological research: estimating biodiversity and providing quantitative estimates of abundance. For both diversity and abundance investigations, we compare estimates across our three simulated community biomass distributions and the three amplification cases at standardized endpoint of 35 PCR cycles.

All of the above simulations and calculations were carried out in R ver 3.5.1 [49] most prominently using packages tidyverse [50] and vegan [51]; all code and related data are available as supplementary material and at https://github.com/invertdna/eDNA_Process_Simulations.

### Biodiversity

#### Effect of PCR Cycle-Number on Sequence Diversity

We examined the effect of the numbers of PCR cycles under three primer efficiency scenarios (Cases A, B, and C above) on over 100 communities of 1000 taxa each, with biomass distributed according to our moderately variable scenario (*γ* = 5). We sampled each community at 5-cycle intervals from 5 to 50 PCR cycles. We estimated sequence diversity using two of the most commonly used metrics of biodiversity, species richness and Shannon diversity. Richness is simply the number of unique taxa identified in the eDNA results, whereas Shannon takes into account both the number of unique taxa as well as their relative frequency.

We note that there is a very large literature examining the measurement and partitioning of diversity [52, 53, 54], and that many different indices have been used to capture the diversity of a community. We include only richness and Shannon diversity here because they are commonly used and they aptly illustrate the issues that arise from using metabarcoding data for studies of biodiversity.

#### Effect of Amplification Bias and Underlying Biomass on Sequence Diversity

To test for the effect of among-taxon amplification bias, we compared biodiversity estimates derived from the three amplification efficiency cases described above and for the three biomass distributions (uniform, low variability, high variability). Each taxon (or equivalently, each unique template molecule) was assigned a fixed amplification efficiency drawn from the case-specific amplification distribution. For all simulations, we compare results after 35 PCR cycles for 100 replicate simulated communities of 1000 taxa each.

#### Quantifying Biomass with Metabarcoding

An aspirational use of eDNA technology is to determine the abundance or biomass of particular species [18, 55, 56, 57, 58]. While research using qPCR or ddPCR technology suggests using single species genetic approaches can yield quantitative estimates of abundance [59, 60, 61], the relationship between amplicon sequence counts and organismal abundance is not straightforward. In particular, single metabarcoded samples in space or time tell us little about the underlying biomass of surveyed organisms, because the amplification efficiencies of each taxon are generally unknown. However, indices of amplicon abundance – reflecting temporal or spatial trends in taxon-specific amplicon abundance – have mirrored biomass in practice [4].

We expect each taxon to have a different amplification efficiency for a given set of PCR primers and therefore expect a poor correlation between eDNA amplicon abundance and biomass abundance when analyzing a dataset of many taxa in a single sample. However, we investigate whether a temporal series of samples can solve this problem; if we assume that amplification efficiency is solely a product of primer-template interaction (and is thus independent of community composition), amplification efficiency remains constant within a taxon across samples. We can then express DNA abundance for each species at each time point as using several alternative metrics (described below) and ask which metrics are likely to be useful for describing the biomass of individual species.

Importantly, this approach relies on the assumption that we need not know a taxon’s efficiency in absolute terms; only that it remains constant across samples. This assumption holds true at least for suites of samples containing identical sets of taxa at different concentrations [4] or samples containing varying subsets of taxa drawn from a common pool [41]. These references show nearly identical within-taxon amplification efficiencies derived from different starting communities: R^2^ = 0.98 (p = 10^−8^, N = 2 communities of 10 fish species at different concentrations using 12s primers; [4]), and median R^2^ = 0.94 and 0.91 (p < 0.01, N = 10 communities of subsets of six fish species drawn from a pool of 10; 12s primers and Cytochrome B primers, respectively; [41]). See Supplementary Information for calculations.

To test the quantitative relationship between biomass and various amplicon-abundance indices, we conducted the simulations described above for 25 time points (spatial points are conceptually equivalent). For each timepoint, we assumed each species randomly varied around a stable abundance and drew a proportional biomass for each species from a symmetric Dirichlet distribution (*γ* = 5) as described above. We then simulated amplifications of each of these communities with a single primer set (Case A, symmetrical) after 35 PCR cycles. To evaluate the performance of a variety of amplicon-based indices, we correlated the biomass of each taxon (N = 1000 in total simulated community, not all of which amplify with the selected primer set) at each time-point (N = 25) against eDNA abundance metrics, reporting the distribution of correlation coefficients (Spearman’s *ρ*) as a summary measure of each index’s quantitative relationship to biomass. We compared each of these to a null distribution derived by randomizing the eDNA amplicon matrix (such that median *ρ* ≈ 0).

Because it is unclear which amplicon summary statistics should be most useful to explain the relationship with biomass, we evaluated a range of indices of amplicon abundance (numbered directly below) against the species specific proportion biomass over the 25 time points. For each equation below, *i* indexes species and *j* indexes sample.

1. Raw amplicon read-counts
2. An index of read-count proportions, scaled 0 to 1 (“eDNA Index”; as used in [4]). Note this is a linear correlate of the *χ*^2^ transformation in [62], of amplicon proportions within a sample, and of the geometric-mean-based adjustment in DESeq2 [63], and so those are not included here. It is also identical to the “Wisconsin double-standardization”, as implemented in vegan [51], with appropriate margins specified. All behave identically.

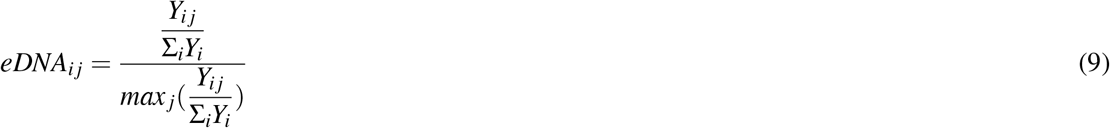
3. Amplicon frequency within a sample, *Freq*, calculated such that the average of non-zero taxa is 1 (method “frequency” in the vegan function “decostand” [51, 64])

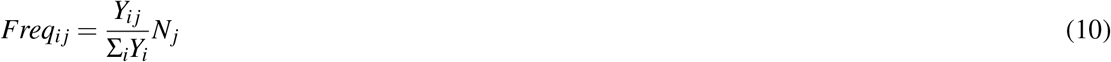
4. Normalized amplicon counts (sample sum-of-squares equal to one)

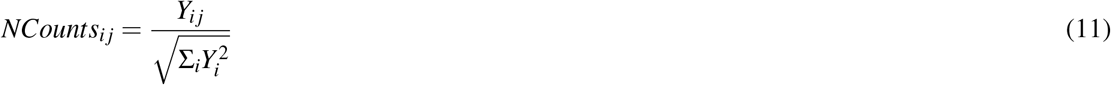
5. Rank order of amplicon abundance, excluding zeros
6. Hellinger distance, a scaled square-root transformation of read counts as defined in [62] and implemented in [51]
7. *log*_2_(*x*) + 1 for values > 0, as implemented in vegan function “decostand”, method “log” [51]

Having measured the performance of these indices by their correlations with simulated taxon biomass, we then decomposed these results to measure the effect of amplicon abundance and amplification efficiency on index performance.

## 3 Results

### 3.1 Diversity Results

#### Effect of PCR Cycles

Our eDNA metabarcoding simulations reveal a strong effect of the number of PCR cycles on estimates of biodiversity (Figure 3A). Increasing the number of PCR cycles decreased both richness and Shannon diversity, but the shape and severity of this decline depended upon the distribution of amplification efficiencies.

**Figure 3.**
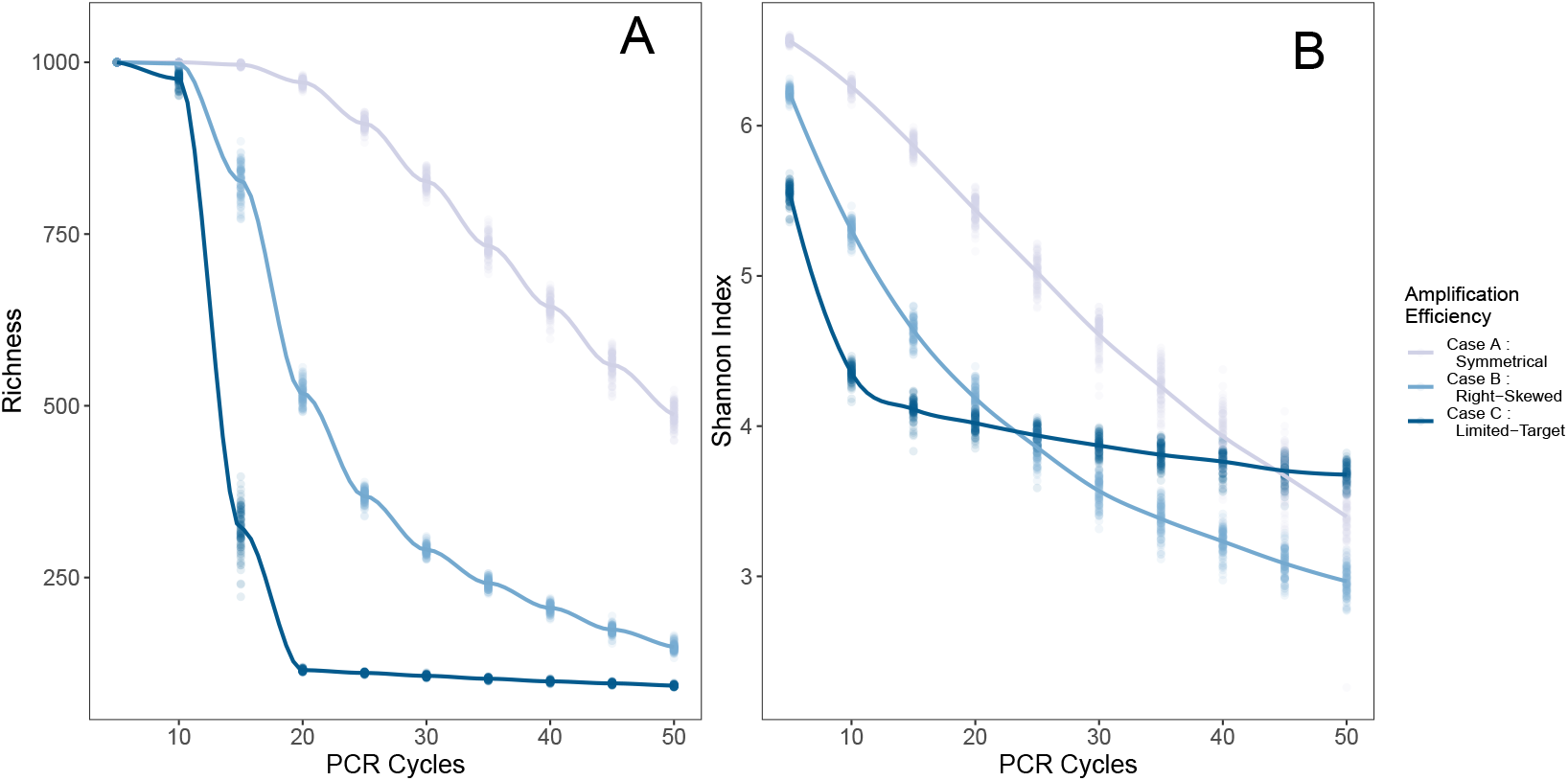
Summary statistics by PCR cycles for 100 simulated communities of 1000 taxa each, keeping amplification efficiency constant for each taxon. Three distributions of amplification efficiencies are shown, color-coded. The underlying biomass distribution is moderately variable (*γ* = 5, as described in Methods. A: richness, B: Shannon Index. Loess-smoothed lines are shown to illustrate trends.

The simulated primer set efficiently amplifying the fewest taxa (Case C) experienced the greatest decline in richness, with a median of only 88 out of 1000 taxa present detected after 20 cycles (N = 100 simulations). This fraction detected mirrors the proportion of taxa amplified with a relative efficiency of greater than approximately 0.6 in the underlying distribution of amplification efficiencies (0.088). By contrast, a primer set that readily amplifies most target taxa (here, Case A, with 63% of the taxa amplifying at efficiency 0.6 or better) predictably recovered the greatest richness, with 973 out of 1000 taxa (median, N = 100) recovered after 20 cycles, and 650 after 40 cycles. Shannon Index values showed similar trends (Figure 3B).

Diversity metrics change rapidly with increasing cycle numbers; for example, estimated richness might fall by half or more between cycle 30 and cycle 40 as in Case B. Such dramatic changes with small analytical differences have two immediate implications: the importance of maintaining consistent procedures within a project (such that results are comparable among samples), and the difficulty of comparing results across datasets generated with even subtly different methods. Furthermore, given that the proportions of eDNA reads are only poorly correlated with the proportions of biomass in most cases (see Results below), the absolute magnitudes of the Shannon Index and similar traditional summary statistics – which depend upon the proportions of each taxon in a community – likely have little meaning in metabarcoding studies.

Case C highlights a notable exception to this idea: a taxon-specific primer, amplifying a small fraction of the total species present, appears stably reflect the richness of the target amplified group after the first few PCR cycles (Figure 3A).

The species-accumulation curves reflect the substantial effect of PCR cycle number of detected richness (Figure 4). These curves illustrate that diversity measures depending upon the slope of species accumulation are themselves strongly influenced by the number of PCR cycles.

**Figure 4.**
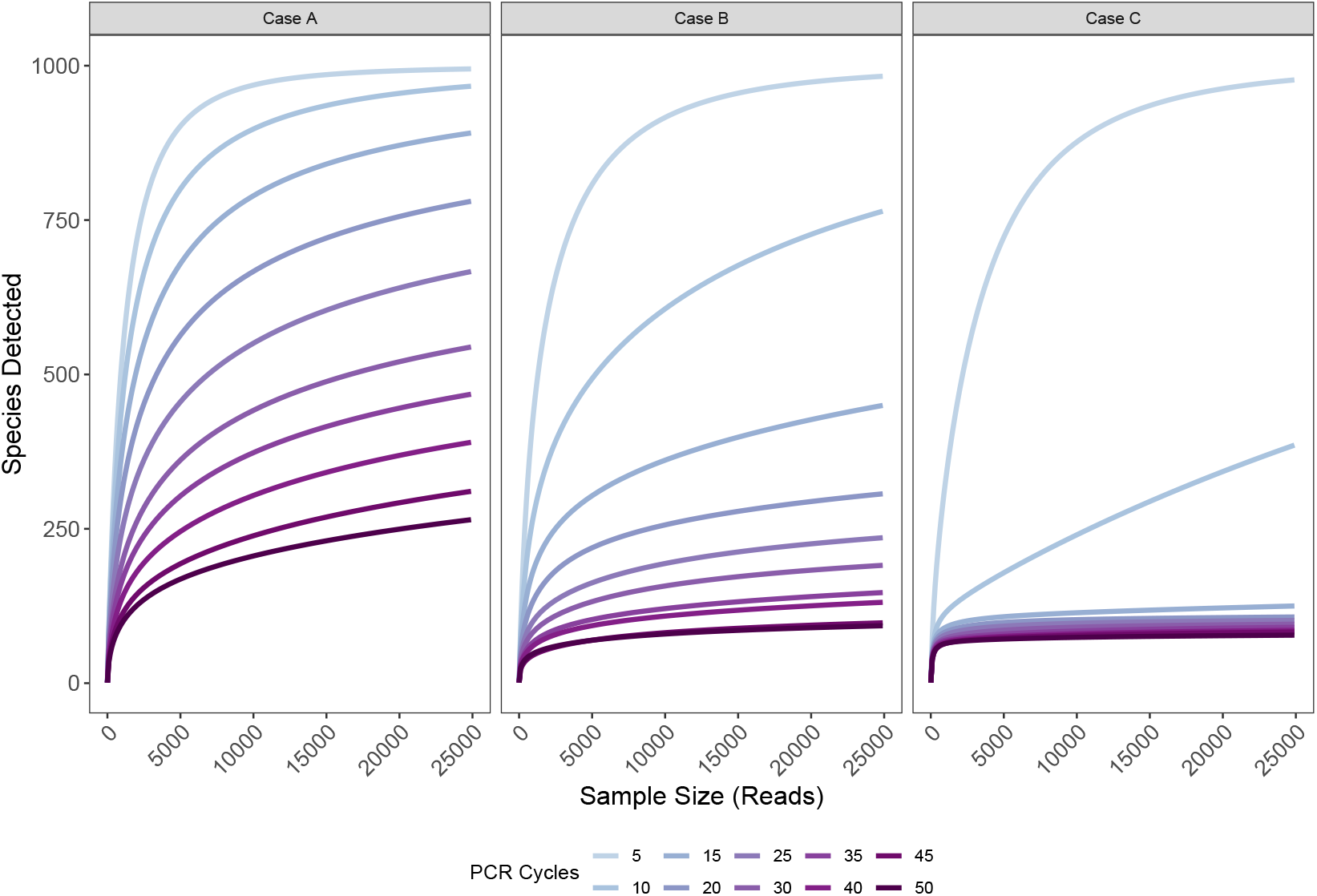
The effect of PCR cycle number on species recovered. Species-accumulation curves for a single simulated community of 1000 target taxa and having moderately-variable proportional biomass, for each set of amplification efficiencies (panels), after simulated sequencing with different numbers of PCR cycles (color).

#### Effect of Amplification Bias

Holding the number of PCR cycles constant – here, for illustration, at 35 cycles – different primer sets yield radically different estimates of diversity in the same simulated communities (Figure 5). More narrowly targeted primer sets predictably reflect lower richness. These findings are consistent with other simulations ([23]) and with empirical results (e.g., [12]), and underscore the broader finding that different primer sets reveal different suites of taxa from a given environment.

**Figure 5.**
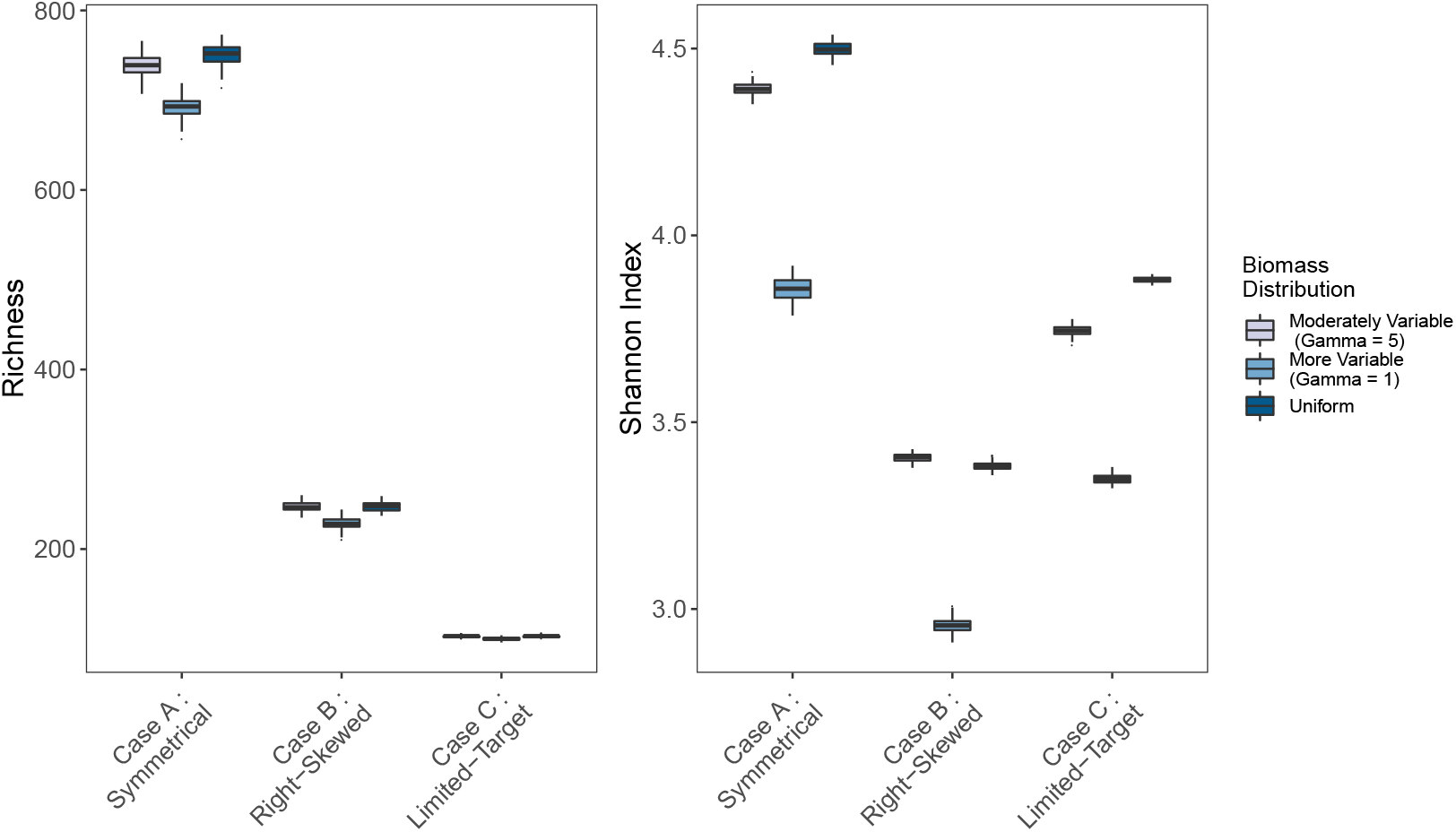
Richness (A) and Shannon Index (B) by PCR cycles for 100 simulated communities of 1000 taxa each after 35 PCR cycles, varying amplification efficiency and varying underlying biomass distributions.

Notably, primer sets performed similarly across quite different distributions of underlying biomass (Figure 5). We can apportion the variance in results attributable to differences in underlying biomass vs. primer-amplification efficiency, keeping the distribution of shedding rates constant, to examine the effects of each. Primer set accounted for more than 99% of the variation in richness, with biomass distribution accounting for far less than 1% (ANOVA; R^2^ = 0.996 and 0.0016, p < 10^−16^ for each). Biomass had a greater influence on Shannon indices, although primer set remained the dominant source of variance (ANOVA; R^2^ = 0.865 and 0.118, p < 10^−16^ for each). These results suggest that metabarcoding results are quite robust to different underlying distributions of biomass, which may or may not be an advantage of the sampling technique, depending on the aims of a particular study.

The probability of detecting any taxon therefore depends upon its amplification efficiency for a given set of primers, and to a much lesser extent, the underlying distribution of biomass or shedding rate (Figure 6). For taxa at a particular amplification efficiency, higher-variance community biomass distributions may lead to higher variance in detectability among taxa. For example, within the median (i.e., fifth) decile bin of amplification efficiency, the variance in likelihood of detection ranged over two orders of magnitude, from 10^−4^ (uniform biomass) to 10^−3^ (moderately variable biomass distribution) to 10^−2^ (more-variable biomass distribution). In sum, communities with greater variability in biomass of target taxa are likely to yield somewhat noisier eDNA datasets, but the qualitative trends appear approximately constant across different biomass distributions.

**Figure 6.**
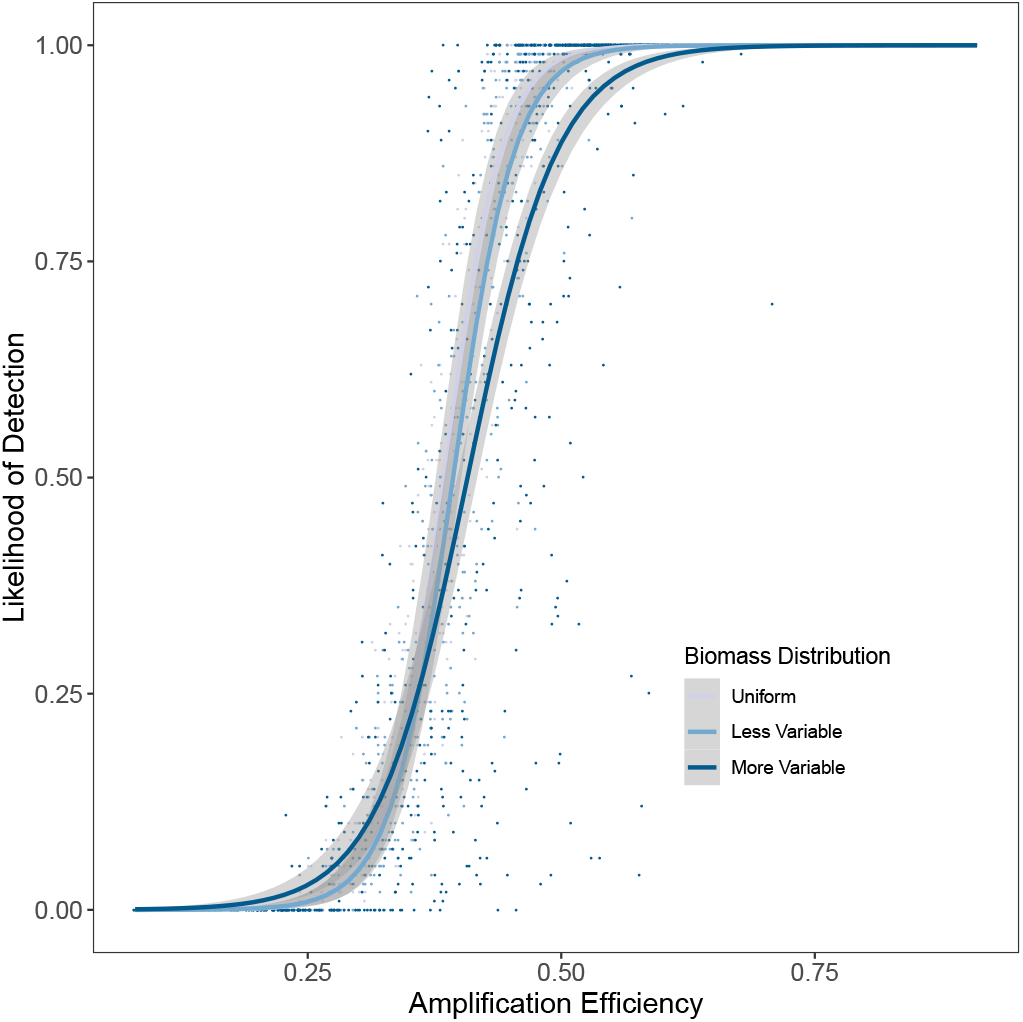
Probability of detection for 1000 simulated taxa after 35 PCR cycles across 100 replicate datasets, as a function of amplification efficiency. The underlying biomass distributions are shown in different colors, and logistic best-fit models added for clarity. The distribution of amplification efficiencies was held constant across datasets (Case A, symmetrical).

#### Quantitative eDNA Indices

Within a given community sample (representing a single timepoint, or equivalently, a single point in space), biomass is only modestly correlated with eDNA abundance (Figure 7; grey vertical lines; median *ρ* = 0.12 - 0.495, biomass vs. different eDNA-abundance indices).

**Figure 7.**
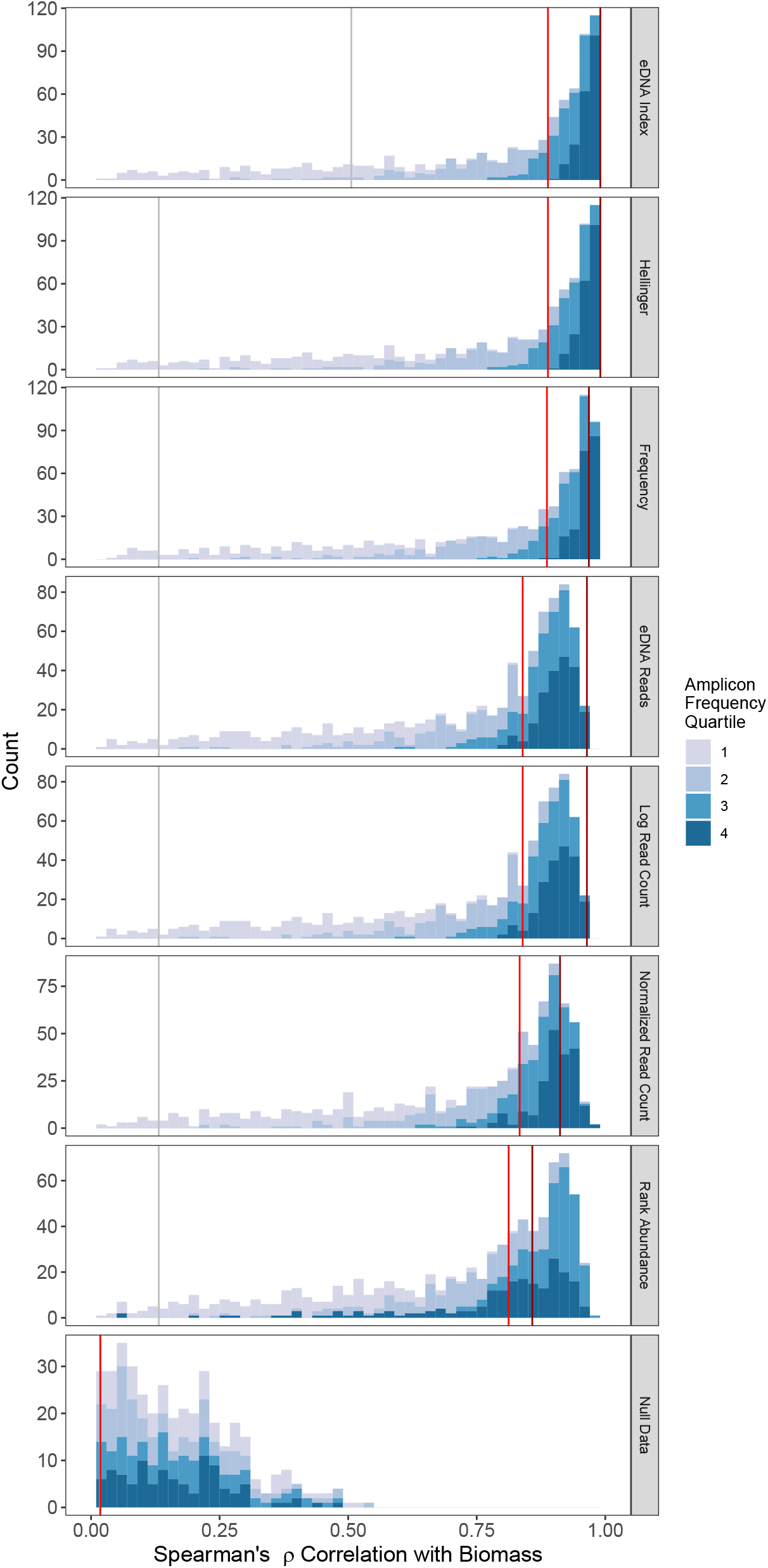
Histograms of Spearman’s rank correlation coefficient (*ρ*), reflecting the relationships between simulated biomass and a variety of eDNA-abundance indices, for a set of 25 simulated time-series samples of a community of 1000 taxa. Shading refers to the quartile of log amplicon frequency; more abundant amplicons are shown in darker shades. Vertical grey lines reflect the median *single-time-point ρ* for that index vs. biomass. Bright red and dark red lines indicate medians and modes, respectively, for the *time-series* indices. Median and mode lines calculated from the underlying data; binning may make maximum values appear different. Correlations calculated for taxa appearing in at least five of the 25 timepoints (i.e., 20% incidence) to avoid many rank ties at zero abundance. The null dataset is the set of correlations between a randomly shuffled amplicon-count matrix and the biomass matrix; this results in a symmetrical distribution of *ρ* with a mean of zero. Only positive values of the null distribution are shown.

When we used the replicate sampling of species across all 25 timepoints, however, many of the indices of eDNA-derived taxon abundance were highly correlated with true biomass (Figure 7). In particular, the index of eDNA-read proportions (“eDNA index”) behaved particularly well, with a median *ρ* of 0.87, and a mode 0.97. For ease of understanding, the eDNA Index is a double-transformation: first, converting amplicon read-counts to proportions (within a sample), and second, scaling the resulting proportions of each read-variant (or OTU, taxon, etc) to the largest observed proportion (across samples) for that read-variant. Various other indices also reliably tracked biomass (Figure 7). All indices perform significantly better than the null expectation derived from the permutation test (Kolmogorov-Smirnoff test, p < 10^−16^). This result suggests metabarcoding studies can indeed reveal detailed information on the abundance of individual taxa.

Taxa with greater amplicon abundances tended to better reflect biomass across all indices investigated (Figure 7, darker shades). For example, for taxa in the first (lowest) quartile of log read abundance, the median eDNA index - biomass *ρ* is 0.4; this rises to *ρ* = 0.77, 0.93, and 0.97 for the second, third, and fourth quartiles respectively. This pattern is likely a function of greater statistical power to detect trends among more-common amplicons, because rare taxa are subject to much greater proportional sampling error.

Moreover, because amplicon abundance depends primarily upon amplification efficiency rather than biomass, the eDNA index almost precisely (median *ρ* = 0.96) tracked taxa with a relative amplification efficiency of greater than approximately 0.6 – regardless of whether their underlying biomass was common or rare (Figure 8). At lower amplification efficiencies, amplicon indexing fails entirely, with the biomass correlation approaching the null distribution when amplification efficiency fell below 0.35 (median *ρ* = 0.09). We suggest the rarity of inefficient amplicons after 35 cycles – combined with process error associated with PCR (ε, in our simulation) and stochastic variability in read-depth – explains this stochasticity.

**Figure 8.**
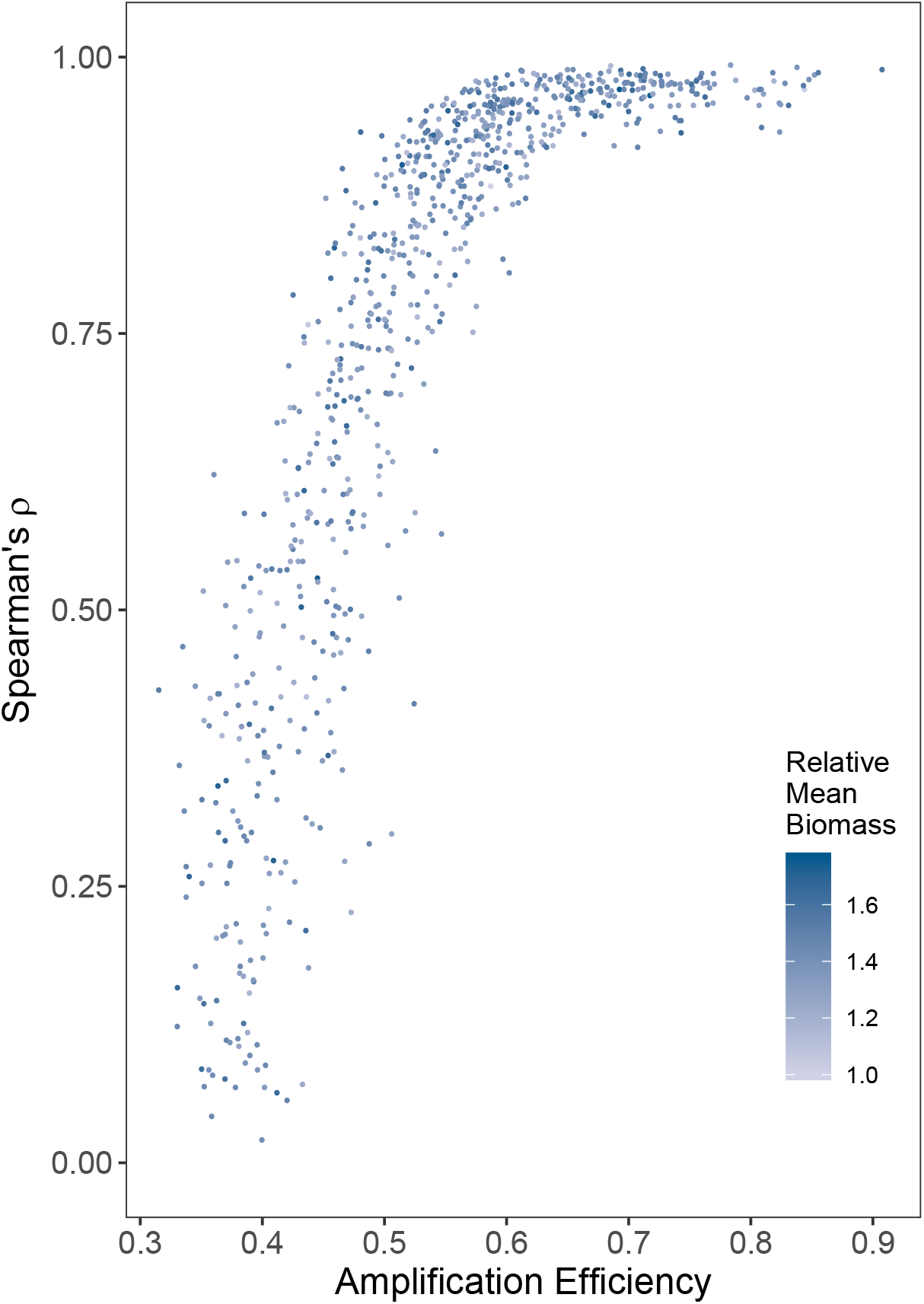
Using the eDNA Index, the biomass-index correlation coefficient (*ρ*) by amplification efficiency for each amplified taxon. Those taxa with a relative amplification efficiency >= 0.6 have particularly strong correlations (median *ρ* = 0.96). As shown by shading, the eDNA Index behaves similarly for species with greater and lesser proportions of biomass in the community. Simulated biomass varied over two orders of magnitude across taxa; averaging across time-points narrows this range to a factor of two, and the relative mean biomass expressed here reflects that smaller range.

Building amplicon indices across different primer sets for the same underlying biological community [65] is a way of creating an ensemble index that can better capture biological dynamics than any single primer set can alone (Supp. Fig 4).

## 4 Discussion

As genetic-based monitoring and discovery tools grow in popularity for ecological applications, it is increasingly important to understand the mechanisms underlying sampling technologies and how these methods affect inferences about ecological communities. We use simulations to identify how two researcher-defined processes in particular – primer choice and the associated amplification distribution, and the number of PCR cycles – can have dramatic consequences for estimates of biodiversity. Additionally, we show how reliable metrics of biomass may be derived from metabarcoding surveys. Together our results help to explain the behavior of PCR-based surveys and suggest clear avenues for integrating eDNA data more fully into ecological applications.

Our simulations suggest three principal conclusions broadly relevant to eDNA work:

1. Traditional ecological diversity metrics – such as richness and the Shannon Index – shift substantially with small changes of PCR-based protocols, to the extent that such metrics may not be comparable across methods or studies. Taxon-specific primer sets are likely to be an exception to this rule because, with a narrow range of target taxa out of the available pool, their results stabilize after a few PCR cycles.
2. The results of community-wide diversity studies depend even more strongly on the choice of PCR primers. Amplification-efficiency explains amplicon abundance to a far greater extent than does underlying biomass within a sample.
3. However, because amplification efficiency is approximately constant for a given taxon and primer set, changes in taxon-specific abundance indices reliably and quantitatively track changes in biomass over space or time. Primer-taxon pairings with relatively high amplification efficiencies are particularly effective in this regard.

We discuss these conclusions in turn below, before suggesting best practices for applying them in the field.

### Exponential Growth: Effect of PCR Cycle Number

Decades after microbial ecologists embraced PCR-based methods [e.g., 66], PCR-based surveys have begun to radically change the way molecular ecologists work with the visible world around them. In mixed-template applications, PCR serves a dual purpose: first, it selects particular DNA fragments of interest; and second, it amplifies these fragments for analysis. In the bargain, however, PCR radically distorts the underlying proportions of biomass as a result of amplification bias [19].

Our simulation shows metabarcoding fails to recover the true value of two traditional biodiversity metrics after as few as 25 PCR cycles. Importantly, the magnitude of difference between the estimated and true values of diversity varies strongly with the distribution of amplification efficiencies, suggesting that results from each combination of primer set and target community will vary unpredictably. And because both exponential amplification and primer bias obscure proportions of species’ biomass, we note that the absolute values of Shannon Index and most other ecological summary statistics – which depend upon species proportions – are likely meaningless in the metabarcoding context.

But measurements of local richness (*α* diversity) and other diversity statistics are rarely studied for just one sample; scientists are often interested on its variation across systems or through an environmental gradient. We find recovered *α* diversity and Shannon Index depend principally on the distribution of amplification efficiencies across the taxa and number of PCR cycles; thus if the same analytical techniques are used consistently, the results will likely accurately reflect relative patterns of diversity.

Similarly, we find after the first few PCR cycles, each cycle greatly magnifies the difference between true and recovered diversity, such that small differences in protocol strongly affect results. Usually PCR protocols are consistent within a project, thus allowing for comparisons between samples processed with a shared protocol, but our results underscore the value of consistent analytical technique. This observation also complicates the prospects for meta-analysis of eDNA-sequencing studies.

### Amplification Bias: Effect of PCR Efficiency

Different PCR primer sets result in vastly different suites of eDNA amplicons [12], an effect described more than twenty years ago in the microbial context [20]. Our simulations suggest the mechanism for such differences is the primer-template interaction, and in particular, the efficiency of amplification: we show – unsurprisingly – that different distributions of amplification efficiencies greatly affect estimates of biodiversity. This result makes clear that metabarcoding studies are not necessarily comparable across systems.

Our simulation suggested amplification-efficiency (i.e., primer bias) had a 630-fold greater impact on richness than did the underlying biomass proportion. This result highlights both a strength and a weakness of eDNA work: depending upon the primer set, the resulting amplicons may at the same time reflect relatively rare taxa and fail to reflect relatively common taxa in a sampled environment.

### Testing Quantitative eDNA Indices

Primer-template bias largely determines the outcome of metabarcoding studies, however, primer-template interaction appears to remain constant across different pools of potential amplicons. As a result, taxon-specific indices constructed from multiple samples taken over time or space appear to quantitatively reflect changes in underlying biomass. Our “eDNA Index” – which, again, is simply an adaptation of transformations that have long existed in ecology – tracks changes in biomass quite closely both in simulations (as here) and in practice (e.g., [4, 59]. Given that many survey applications demand a degree of quantification, we view this as an important finding. Nevertheless, we note that a quantitative index is not the same thing as counting actual target species. Tying the changes in an eDNA index to an actual number of individuals of a species (or kilograms of biomass), for example, will likely require calibrating the index against samples of known composition in a field setting.

### Best Practices

Mindful of the recommendations contained in series of existing review papers on eDNA [67, 68, 69], we offer the following suggestions for standardizing eDNA techniques in light of our own findings.

- To maximize diversity detected with a given primer set, minimize PCR cycles, preferably fewer than 35.
- Keep PCR protocols strictly consistent across samples you wish to compare.
- Do not compare absolute values of richness, Shannon Index, or similar metrics across studies.
- Be specific about a target organismal or ecological group before sampling, in order to define the species expected and a denominator for total expected diversity. This may take iteration and experience with a particular primer set.
- For each primer set, estimate the distribution of amplification efficiencies within your target group using mock communities or other calibration techniques. This will set an expectation for the fraction of target taxa recovered and define amplification bias among the recovered species.
- Carry out a temporal or spatial series of samples in order to track organismal changes using an index of eDNA abundance.

## 5 Conclusion

The results of metabarcoding studies differ dramatically from those of traditional, non-PCR-based sampling methods as a result of the PCR process itself. This exponential process means that 1) small changes in laboratory technique can yield large differences in outcomes, 2) PCR-based assays likely act differently on every target species, 3) there is consequently no one-to-one correspondence between the number of assigned reads in an eDNA study and the abundance of the source organism, and 4) neither might we expect a universally strong correlation in estimates of taxon-richness between eDNA and traditional methods.

Nevertheless, the power of metabarcoding surveys is undeniable: the technique reveals hundreds or thousands of taxa in every sample, and can easily distinguish ecological communities among habitats and sampling sites. Many practical applications demand some quantification of organisms – for example, fisheries stock assessments, or population surveys for endangered species – and so understanding the processes linking amplicon reads to species’ biomass or counts is particularly relevant for making eDNA a standard source of data for ecological sampling. By focusing on the processes by which metabarcoding results arise, we have developed a picture of the specific ways in which these might – and might not – be compared to other survey techniques, and in the process, have provided a quantitative means of tracking changes in environmental samples.

We note that our results are consistent with [22] – a draft of which became available at approximately the same time as our original manuscript submission – which treats quite similar subject matter from a statistical, rather than molecular biological, perspective. Taken together, along with other recent work such as [23] and [21], a common understanding of the processes underlying PCR-based studies appears to be coalescing.

## 5.1 Author Contributions

RG conceived the project, edited the manuscript, and provided routine feedback during development. AOS contributed significant philosophical and statistical expertise, edited the manuscript, drafted some of the final text, and consulted during project development. RPK carried out the main analyses, wrote the main manuscript text, and prepared the figures. All authors reviewed the manuscript.

## 5.2 Acknowledgements

We are grateful to M. Stoeckle, J. Ausubel, and the other organizers of the 2018 National Conference on Marine Environmental DNA at Rockefeller University for providing us an opportunity to think through these issues. We thank Emily Jacobs-Palmer for thoughtful input and consistent support. We thank K. Cribari for lab assistance, as well as the UW Center for Environmental Genomics and Linda Park’s lab at NOAA Fisheries. The reviews of an editor and (especially) one anonymous reviewer were very helpful in improving the manuscript. A version of this manuscript appeared on bioRxiv.org (https://doi.org/10.1101/660530) under its pre-peer-review title, “Understanding Environmental DNA.”

## 5.3 Additional Information

The author(s) declare no competing interests.

## 5.4 Data Availability

All materials, analytical code, and resulting data are available as supplementary files accompanying this manuscript, as well as on GitHub at https://github.com/invertdna/eDNA_Process_Simulations.

