## Supplemental_info for "Understanding PCR Processes to Draw Meaningful Conclusions from Environmental DNA Studies"

### Supplementary Information

#### Empirical Data on Amplification Efficiency Distributions Among Species

| Reference | Gene Region | Primer Name | Target Group | Sample Size | Shape 1 ± SD | Shape 2 ± SD |
|---|---|---|---|---|---|---|
| Port et al. 2016 | 12s | Riaz 12s | Fish | 10 | 1.54 ± 0.44 | 0.61 ± 0.12 |
| Hänfling et al. 2016 | 12s | Riaz 12s | Fish | 22 | 2.1 ± 0.54 | 0.58 ± 0.1 |
| Hänfling et al. 2016 | Cytochrome B | L14841/H15149 | Fish | 22 | 0.97 ± 0.19 | 0.65 ± 0.11 |
| Olds et al. 2016 | Cytochrome B | L14841/H15149 | Fish | 6 | 0.52 ± 0.11 | 0.56 ± 0.12 |
| Olds et al. 2016 | 12s | Am12s | Fish | 6 | 0.42 ± 0.08 | 0.64 ± 0.14 |
| Olds et al. 2016 | 16s | Ac16s | Fish | 6 | 1.24 ± 0.34 | 0.58 ± 0.12 |
| Olds et al. 2016 | 12s | Ac12s | Fish | 6 | 0.51 ± 0.1 | 0.59 ± 0.13 |
| Deiner et al. 2016 | COI | Folmer | Eukaryotes | 33 | 0.72 ± 0.12 | 0.81 ± 0.14 |
| Ford et al. 2016 | 16s | Salmon-F/16s-R | Fish | 5 | 1.35 ± 0.4 | 0.55 ± 0.12 |
| Braukmann et al 2019 | COI | C_LepFolF/C_LepFolR | Arthropods | 374 | 1.72 ± 0.12 | 1.23 ± 0.08 |

**Supplementary Table 1**: Empirical datasets from which the distribution of amplification efficiencies for primers and target taxa could be calculated. Each paper provided its own data (in supplementary information of each of the original references) for one or more mock communities of known composition and DNA concentration from a variety of species. Here, we show the number of species tested in the original papers as "Sample Size". We used the equation $A_i = D_i(a_i + 1)^{N_{PCR}}$, as given in the main text, to derive species-specific values for amplification efficiency, $a_i$. We then fit a Beta distribution using a Hamiltonian Monte Carlo search algorithm, implemented in rstan [1, 2], to each of the sets of efficiencies (one per original paper), and we report the resulting mean shape parameters and standard deviations here. We note that degenerate primers (as in Deiner et al. 2016) and cocktails of similar primers (as in Braukmann et al. 2019) are essentially the same thing in practice, and so we include both here.

Two references provided results for multiple mock communities, allowing us to assess the consistency of our derived species-specific values of amplification efficiency, $a_i$, in the context of different pools of potential template molecules.

Port et al. 2016 [8] included two communities of 10 fish species at different concentrations using 12s primers (Supplementary Fig. 1). Hänfling et al. 2016 [9] tested 10 communities of six fish species each, drawn from a common pool of 10, using both 12s primers and Cyt B primers (Supplementary Figs. 2 and 3). Thus, the two papers provide a fair amount of data about the consistency of template-primer interaction in different contexts. These references show nearly identical within-taxon amplification efficiencies derived from different starting communities: $R^2 = 0.98$ ($p = 10^{-8}$, N = 2 communities of 10 fish species at different concentrations using 12s primers; [8]), and median $R^2 = 0.94$ and 0.91 ($p < 0.01$, N = 10 communities of subsets of six fish species drawn from a pool of 10; 12s primers and Cyt B primers, respectively; [9]).

#### Building an Ensemble eDNA Index Across Primer Sets

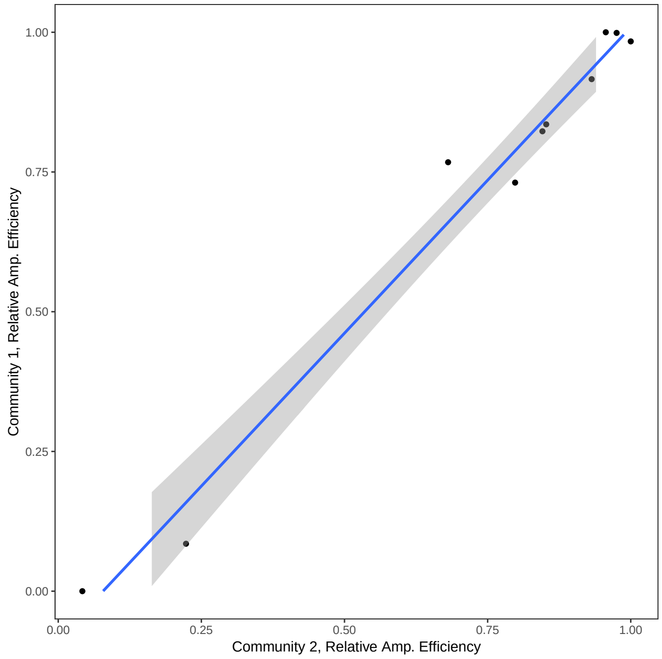


**Supplementary Figure 1**: Best-fit line ($R^2$ = 0.98) for the relative amplification efficiency for 10 species in two mock communities of different mixes of species' DNA concentrations, using 12s primers. Data from [8].


**2.** McElreath, R. *rethinking: Statistical Rethinking book package* (2016). R package version 1.59.

**3.** Riaz, T. *et al.* ecoPrimers: inference of new DNA barcode markers from whole genome sequence analysis. *Nucleic Acids Res.* **39**, e145–e145 (2011). URL http://nar.oxfordjournals.org/content/39/21/e145.short.

**4.** Olds, B. P. *et al.* Estimating species richness using environmental DNA. *Ecol. Evol.* **6**, 4214–4226 (2016).

**5.** Deiner, K., Fronhofer, E. A., Mächler, E., Walser, J.-C. & Altermatt, F. Environmental DNA reveals that rivers are conveyer belts of biodiversity information. *Nat. communications* **7**, 12544 (2016).

**6.** Ford, M. J. *et al.* Estimation of a killer whale (orcinus orca) population's diet using sequencing analysis of DNA from feces. *Plos One* **11**, e0144956 (2016).

**7.** Braukmann, T. W. A. *et al.* Metabarcoding a diverse arthropod mock community. *Mol. Ecol. Resour.* **0**. URL https://onlinelibrary.wiley.com/doi/abs/10.1111/1755-0998.13008. DOI 10.1111/1755-0998.13008. https://onlinelibrary.wiley.com/doi/pdf/10.1111/1755-0998.13008.

**8.** Port, J. A. *et al.* Assessing vertebrate biodiversity in a kelp forest ecosystem using environmental DNA. *Mol. Ecol.* **25**, 527–541 (2016).

**9.** Hänfling, B. *et al.* Environmental DNA metabarcoding of lake fish communities reflects long-term data from established survey methods. *Mol. Ecol.* **25**, 3101–3119 (2016).

**10.** Schloerke, B. *et al.* *GGally: Extension to 'ggplot2'* (2018). URL https://CRAN.R-project.org/package=GGally. R package version 1.4.0.

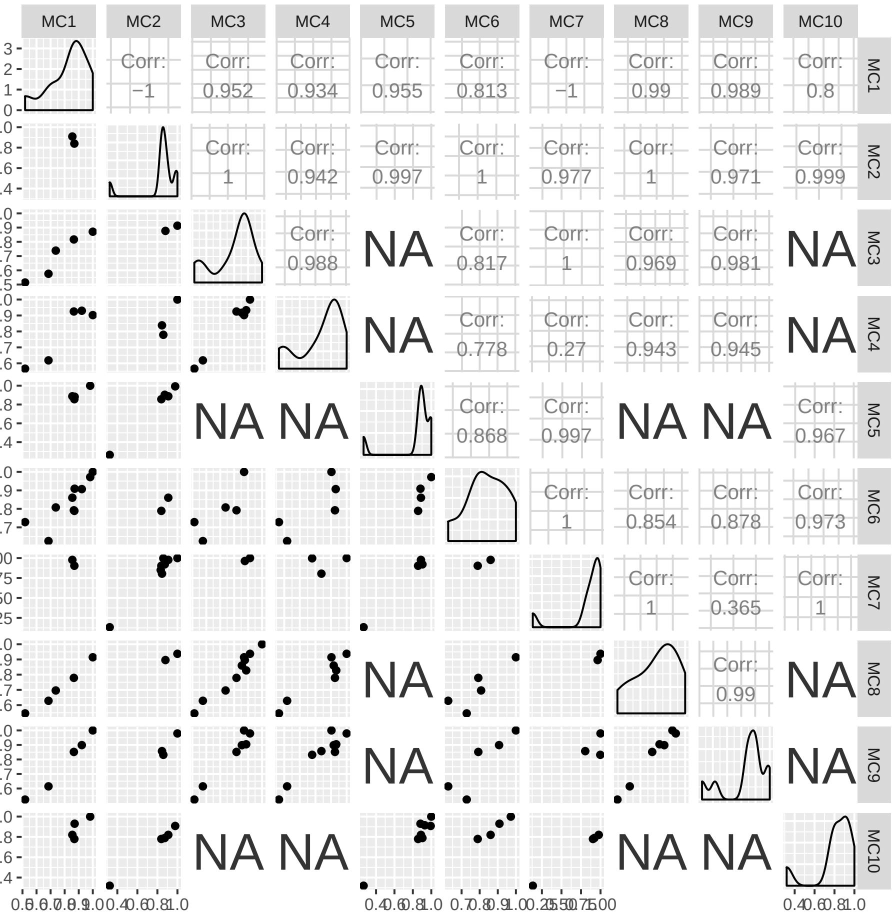


**Supplementary Figure 2**: Paired plots and correlation coefficients for 10 mock communities using 12s primers in [9]. Figure made with [10].

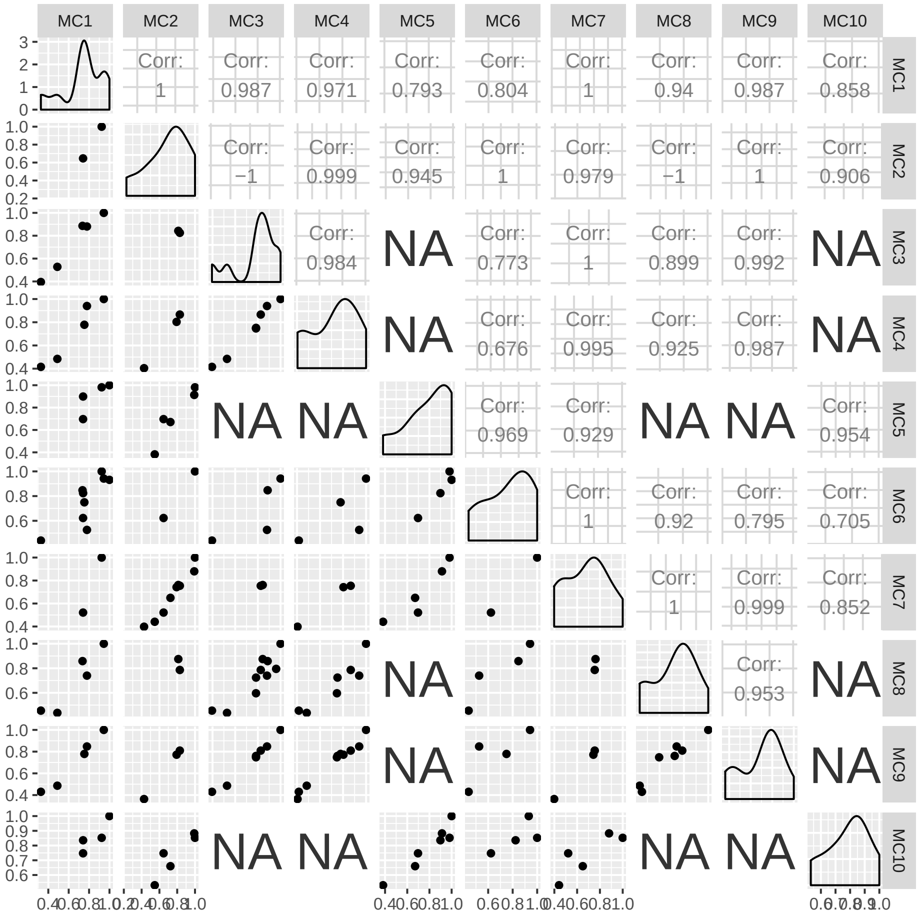


**Supplementary Figure 3**: Paired plots and correlation coefficients for 10 mock communities using CytB primers in [9]. Figure made with [10].

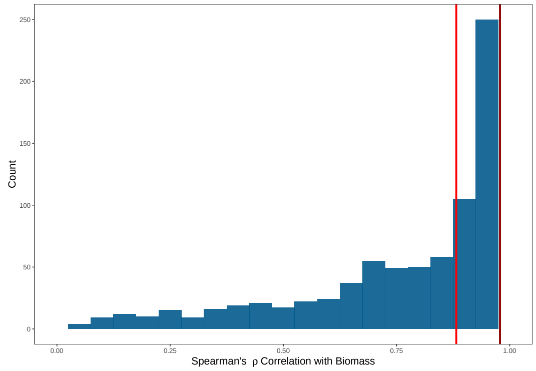


**Supplementary Figure 4**: Distribution of Spearman's rho ($\rho$) for eDNA Index vs. Biomass using a three-locus ensemble index for the 25-timepoint simulation as described in the main text. Of the 1000 species in the simulated community, 879 are amplified by at least one of the three primer sets, in contrast to the single-locus simulations in the main text, which amplify a median 89 - 730 taxa after 35 PCR cycles in our simulations. Thus the ensemble has the advantages of 1) quantitatively combining information across primer sets, and 2) maximizing the diversity of taxa surveyed, while 3) maintaining a strong correlation with changes in biomass. Code for combining loci into an ensemble is included in the supplementary Rmarkdown file that contains all of the analytical code for the main paper.
